# Glucose extends lifespan through enhanced intestinal barrier integrity in *Drosophila*

**DOI:** 10.1101/2020.03.20.000968

**Authors:** Anthony Galenza, Edan Foley

**Author notes:** Corresponding Author: Edan Foley.

## Abstract

Dietary intervention has received considerable attention as an approach to extend lifespan and improve aging. However, questions remain regarding optimal dietary regime and underlying mechanism of lifespan extension. Here, we asked how glucose-enriched food extends the lifespan of *Drosophila*. We showed that glucose-dependent lifespan extension is independent of caloric restriction, or insulin activity, two established mechanisms of lifespan extension. Instead, we found that flies raised on glucose-enriched food increased the expression of cell junction proteins, and extended intestinal barrier integrity with age. Furthermore, chemical disruption of the intestinal barrier removed the lifespan extension associated with glucose-treatment, suggesting that glucose-enriched food prolongs adult viability by enhancing the intestinal barrier. We believe our data contribute to our understanding of intestinal health and may help efforts to develop preventative measures to limit the effects of aging and disease.

## INTRODUCTION

Increases in longevity require parallel interventions that extend healthspan. In this context, nutrition optimization to promote healthy aging has received considerable attention (Kalache et al. 2019). Nutritional deficiencies increase risks of a range of age-related chronic diseases, but we have limited knowledge of dietary interventions that extend life and healthspan (Shlisky et al. 2017). The most thoroughly studied dietary intervention, caloric restriction, extends lifespan in several vertebrate and invertebrate models, though the implications for humans are unclear (Most et al. 2017). Further studies generated interest in protein restriction or intermittent fasting to extend lifespan (Fontana & Partridge 2015). Recent focus has shifted to nutritional geometry, or the effect of mixtures of dietary components rather than isolated nutrients (Simpson & Raubenheimer 2009; Solon-Biet et al. 2015). These studies revealed that low protein to carbohydrate ratios extend longevity of mice and flies, with maximal benefits for a 1:16 protein:carbohydrate ratio in flies (Lee et al. 2008; Solon-Biet et al. 2014). However, the direct effect of carbohydrates on health and longevity are complex, and require further investigation (Lee et al. 2015).

High sugar intake has long been considered a health hazard (Kroemer et al. 2018), particularly in association with obesity, Type 2 diabetes, and cardiometabolic risk (Prinz 2019), and model systems are commonly used to study effects of sugar on health and longevity. For example, providing *C. elegans* 5-50 mM glucose shortens lifespan (Schulz et al. 2007; Schlotterer et al. 2009). Interestingly, 111 mM glucose treatment in young worms (1-3 days old) reduces lifespan, but beginning glucose treatment after worms are at a post-reproductive age (7 days old) extends lifespan (Lei et al. 2018).

The vinegar fly, *Drosophila melanogaster*, is a useful model to study the effects of sugar on aging for a number of reasons. A completely defined holidic medium allows for manipulation of individual ingredients, while allowing for easy comparison between labs (Piper et al. 2014). Major nutrient-sensing pathways, such as insulin and TOR signaling, are highly conserved between flies and mammals and techniques for studying aging in flies are well-established (Piper & Partridge 2017). High-sucrose treatment (1.0 M compared to 0.15 M controls) reduces *Drosophila* lifespan (Na et al. 2013), even with transient exposure (1.2 M compared to 0.15 M controls) in young adults (Dobson et al. 2017). The type of sugar is important, as sucrose is more detrimental than glucose or fructose (Lushchak et al. 2014). Recently, we found that the addition of 0.56M glucose to the holidic medium that contains 0.05M sucrose extends lifespan in *Drosophila* compared to the unmodified holidic medium (Galenza et al. 2016).

In this study, we found that glucose supplementation extends lifespan independent of caloric restriction, or effects on insulin pathway activity. Instead, we showed that glucose-enriched food extends the lifespan of adult *Drosophila* by improving the maintenance of intestinal barrier integrity in aging flies. Glucose-treated flies have increased expression of cell junction proteins and higher levels of the septate junction protein Coracle localized at intestinal cell membranes, and flies raised on glucose-enriched food maintain barrier function to a later age than their control counterparts. Combined, these data presented here identify a relatively uncharacterized diet-dependent mechanism of lifespan extension.

## RESULTS

### Glucose-enriched food promotes maintenance of energy stores with age

In a longitudinal study of relationships between nutrition, age, and metabolism, we found that glucose-enriched (100 g/L) holidic food extends the lifespan of adult *Drosophila* compared to unmodified holidic food, particularly in males (Galenza et al. 2016). As prolonged consumption of sugar-rich food is typically associated with diminished health and lifespan outcomes, we asked how addition of glucose extends longevity for flies. Before addressing this question, we first tested a range of glucose concentrations to identify the optimal amount required for increased longevity. Specifically, we measured longevity of flies raised on holidic food that we supplemented with 0 to 200 g/L glucose. We found that addition of 50 g/L glucose had the greatest effect, leading to a 27% increase in median lifespan compared to unmodified food (Figure S1). Thus, for the remainder of this study we determined the effects of holidic food (HF), and 50 g/L glucose-enriched holidic food (GEF) on health and longevity.

First, we asked what impact added glucose has on metabolism by comparing macronutrient content in wild-type flies raised on HF or GEF for 20 or 40 days. We found no difference in weight either at day 20 or 40 (Figure 1a). Likewise, protein levels remained comparable between flies raised on HF or GEF at both times (Figure 1b). When we measured total glucose, we found that flies raised on GEF had higher levels of glucose at day 40 (Figure 1c). Similarly, we found that triglyceride levels were significantly higher at day 40 in GEF-treated flies than in HF-treated flies (Figure 1d). As GEF elevated total glucose content, we asked if GEF also impacted levels of circulating glucose and trehalose, the primary blood sugar in insects. We found that flies raised on GEF had elevated total circulating sugars at day 40 compared to flies raised on HF (Figure 1e). Looking at the component circulating sugars, this difference is attributable to increased free glucose (Figure 1f), with no detectable effects on trehalose (Figure 1g). As GEF primarily imapcts macronutrient levels in older flies, we used linear regression analysis to ask if GEF affects age-dependent changes in macronutrients. Although not statistically significant, we found that GEF had modest attenuating effects on age-dependent declines in total and free glucose (Figure 1h), and significant attenuating effects on age-dependent loss of triglyceride (Figure 1h). Combined, our data suggest that increased provision of glucose contributes to the maintenance of energy-rich triglycerides and sugars as flies age.

**Figure 1.**
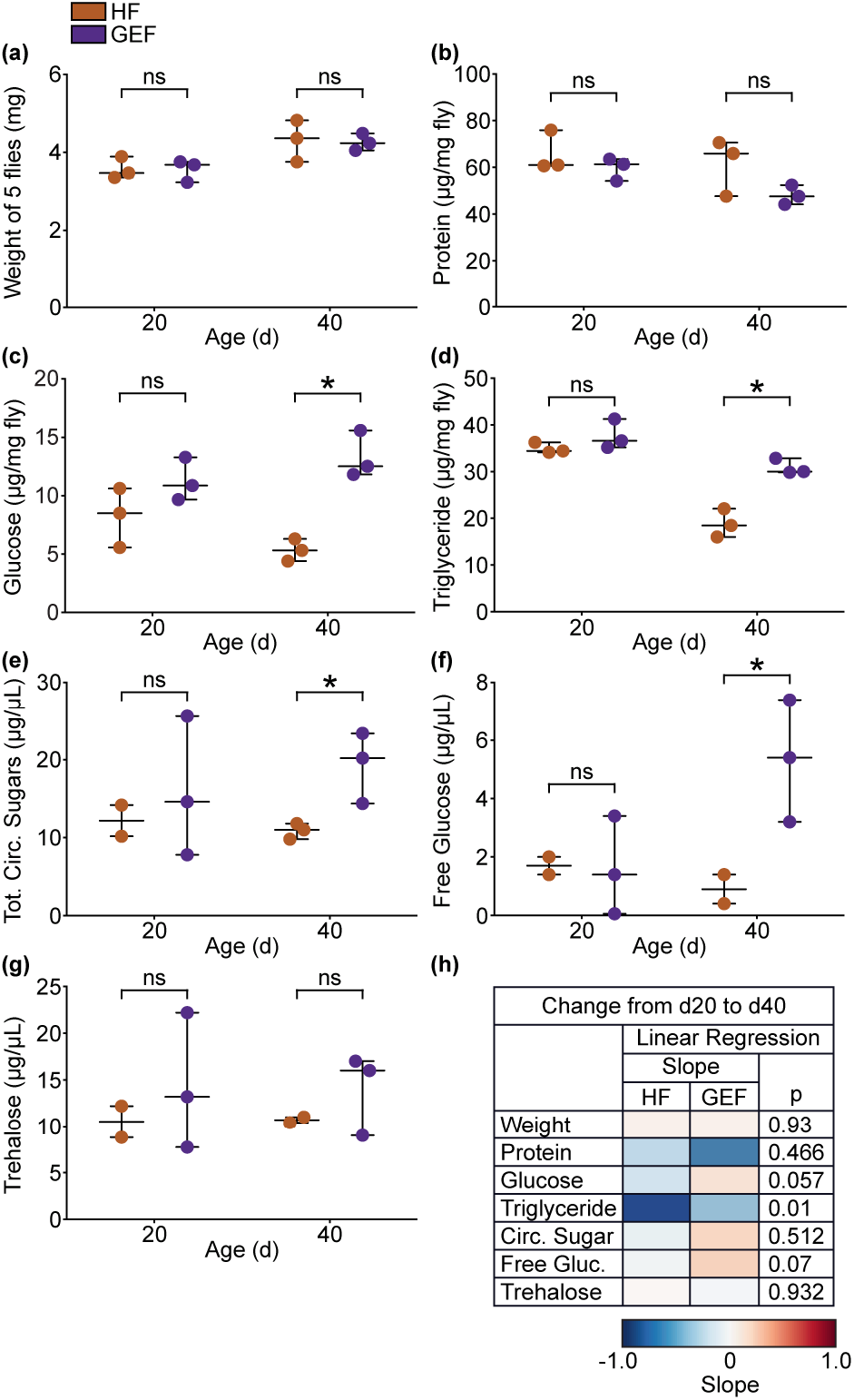
Glucose-enriched food promotes maintenance of macronutrients with age. **(a-d)** Quantification of (a) weight, (b) protein, (c) glucose, and (d) triglycerides in *w^1118^* flies raised on glucose-enriched food (GEF) versus unmodified holidic food (HF) for 20 or 40 days (n = 3). Each dot represents 5 flies. **(e-g)** Quantification of (e) total circulating sugars, (f) free glucose, and (g) trehalose *w^1118^* flies raised on GEF or HF for 20 or 40 days (n = 2-3). Statistical significance (denoted by asterisk) for (a-g) determined by Student’s T-test (p < 0.05). **(h)** Linear regression analysis of data shown in (a-g).

### Glucose-enriched food increases calorie intake

As our flies are fed *ad libitum*, we do not know if GEF-dependent effects on macronutrients are an indirect result of changes in feeding. We consider this an important question to address, as calorie intake and feeding frequency are linked to lifespan in several experimental organisms (Fontana & Partridge 2015).

To measure feeding frequency, we used the flyPAD to count individual sips; bursts, which are clusters of sips; and bouts, which are clusters of bursts, in flies raised on HF or GEF. For this assay, we raised flies on their respective foods for 20 days, then starved them for 2 hours prior to feeding in a flyPAD arena for 1 hour. We saw no difference in sips (Figure 2a), bursts (Figure 2b), or bouts (Figure 2c), between flies raised on HF or GEF, suggesting that GEF does not significantly alter feeding behavior over short periods. To determine if GEF impacts feeding behavior over longer timeframes, we used the capillary feeding (CAFE) assay, to calculate food consumption across three days. In the CAFE assay, flies are fed through capillary tubes that allow us to quantify liquid food consumption. We raised flies on HF or GEF for 20 days before transfer to the CAFE setup for a 3-day period, where flies were fed a liquid version of their respective food. We found that flies raised on HF consumed a greater volume than those raised on GEF, about a 1.2-fold daily increase (Figure 2d). This translates to a 2.3-fold increase in calorie intake for GEF-treated flies compared to HF-treated (Figure 2e). The increased calorie intake is a result of elevated carbohydrate consumption, as flies raised on GEF consumed approximately 3.2-fold more calories from carbohydrates per day than their counterparts raised on HF (Figure 2f). Conversely, amino acids provided approximately 20% fewer calories to flies raised on GEF than on HF (Figure 2g). Together, these data indicate that flies raised on GEF are not calorically restricted, in fact, they consume significantly more calories in the form of carbohydrate. To test if the lifespan extension observed for flies raised on GEF is simply a consequence of feeding adults a higher calorie food, we measured the lifespans of flies raised on modified holidic food isocaloric to GEF, where extra energy was provided either from lard, or protein. As expected, flies raised on GEF lived significantly longer than counterparts on HF (Figure 2h). In contrast, protein-supplemented holidic food had no detectable effects on lifespan, whereas lard-supplemented holidic food shortened lifespan, and significantly increased the risk of early death (Figure 2i). Thus, simply adding extra calories to HF does not extend longevity, indicating that GEF extends lifespan through a more specific mechanism.

**Figure 2.**
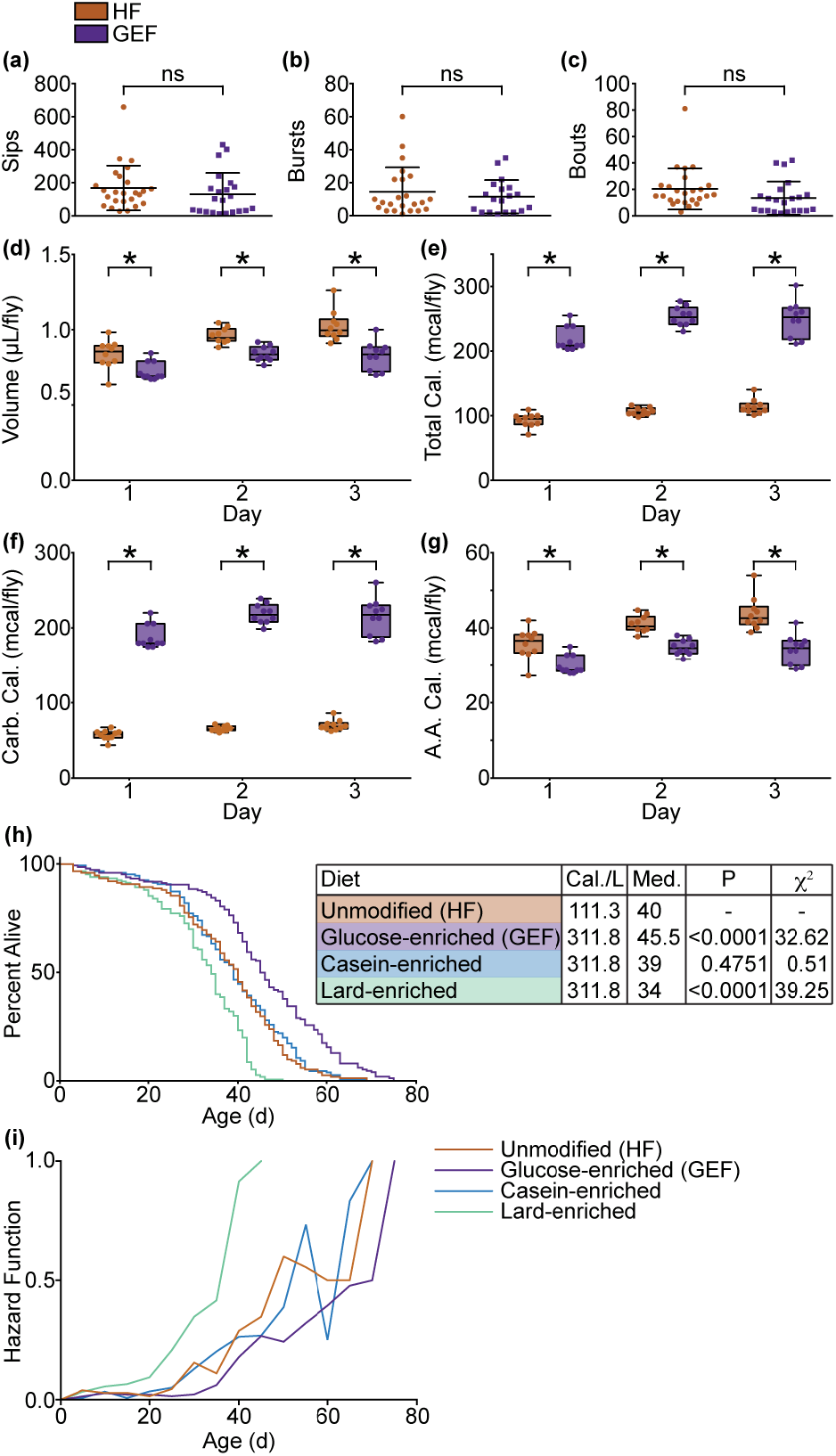
Glucose-enriched food increases caloric intake. **(a-c)** Quantification of (a) number of sips, (b) feeding bursts, and (c) feeding bouts in 20-day old *w^1118^* flies raised on glucose-enriched food (GEF) or unmodified holidic food (HF) using a flyPAD. **(d-g)** Quantification of liquid food consumption in 20-day old *w^1118^* flies raised on GEF versus HF using a CAFE measuring (d) volume consumed, (e) total calories, (f) calories from carbohydrates, and (g) calories from amino acids (AA). **(h-i)** (h) Survival curve and (i) hazard function of *w^1118^* flies raised on HF, GEF, casein-enriched food, or lard-enriched food. (h) Statistical significance determined by log-rank (Mantel-Cox) test shown in table.

### Glucose-enriched food extends lifespan independent of insulin activity

As we observed increased total and circulating glucose in flies that we raised on GEF, we wondered what effects GEF has on the insulin pathway, a known modifier of longevity.

To answer this question, we quantified transcription of the insulin-like peptides (ILP) *ilp2*, *ilp3*, and *ilp5*, in flies raised on HF or GEF for 20 or 40 days. We found that expression of *ilp2* and *ilp5* was lower in 40-day old flies raised on GEF compared to flies raised on HF (Figure 3a, c), while the expression of *ilp3* was unaffected (Figure 3b). In flies, *ilp* gene expression is complex, and does not necessarily reflect amount of peptide in storage, or in circulation (Park et al. 2014). Thus, we used an ELISA to quantify total, and circulating amounts of FLAG and HA epitope-tagged ILP2 (ILP2-FH) in flies raised on GEF or HF. In this line, ILP2-FH expression is controlled by the *ilp2* promoter, and accurately reports ILP2 protein levels (Park et al. 2014). We observed significantly lower total amounts of ILP2-FH in GEF-treated flies compared to age-matched HF-treated controls (Figure 3d). However, we did not detect food-specific effects on levels of circulating ILP2-FH (Figure 3e).

**Figure 3.**
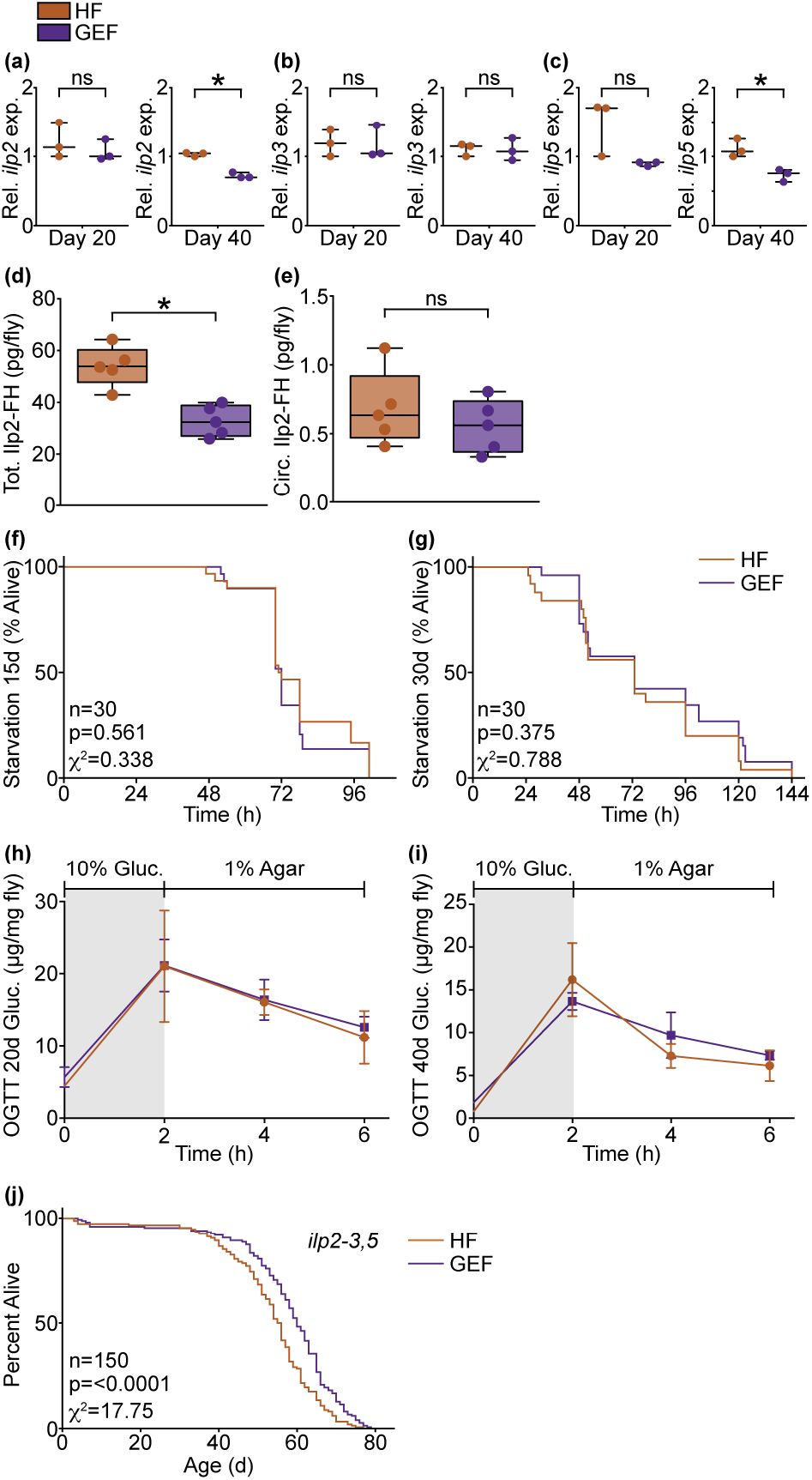
Glucose-enriched food extends lifespan independent of insulin activity. **(a-c)** Quantification of the relative expression of (a) *ilp2*, (b) *ilp3*, and (c) *ilp5* in *w^1118^* flies raised on glucose-enriched food (GEF) versus unmodified holidic food (HF) for 20 or 40 days. **(d-e)** Quantification of (d) total and (e) circulating Ilp2-FH in *w^1118^* flies raised on GEF versus HF for 20 days. Statistical significance (denoted by asterisk) for (a-e) determined by Student’s T-test (p < 0.05). **(f-g)** Survival curve upon starvation of *w^1118^* flies raised on GEF versus HF for (f) 15 or (g) 30 days. **(h-i)** Oral glucose tolerance test performed on *w^1118^* flies raised on GEF versus HF for (h) 20 or (i) 40 days. **(j)** Survival curve of *ilp2-3,5* flies raised on GEF versus HF. Statistical significance for survival curves determined by log-rank (Mantel-Cox) test.

To determine if GEF-dependent shifts in insulin peptide expression translate into effects on insulin activity, we measured starvation resistance and oral glucose tolerance in flies raised on GEF and HF. In flies, insulin impairs starvation resistance, and improves glucose tolerance. Thus, we expect that any effects of GEF on insulin signaling will have measurable impacts on starvation resistance or glucose tolerance. For starvation assays, we raised flies on HF, or GEF, for 15 or 30 days, and followed survival after switching to nutrient-deficient medium. For both ages, we did not detect food-dependent effects on starvation resistance (Figure 3f, g). For the oral glucose tolerance test (OGTT) we raised flies on HF or GEF for 20 or 40 days, followed by a 16h fast, prior to a 2h *ad libitum* feed on a 10% glucose medium, followed by a period of re-fasting. We quantified total glucose in flies following the initial fast (0h), after feeding on 10% glucose (2h), and twice during the re-fast period (4h, 6h). In insulin-sensitive flies, glucose levels rise during feeding, and drop during the fast, due to insulin-dependent stimulation of glucose uptake. We found that flies raised on either food processed glucose with equal efficiency at all time points in both ages (Figure 3h, i), arguing that GEF does not significantly impair insulin sensitivity as the flies age.

Finally, we measured the lifespans of HF and GEF-treated *ilp2-3*, *5* mutant flies. These mutants are deficient for insulin signaling, and outlive wild-type controls. Thus, if insulin signaling is required for GEF-mediated extension of lifespan, we predict that *ilp2-3*, *5* mutants will not benefit from a lifelong culture on GEF. Contrary to our hypothesis, *ilp2-3*, *5* mutants raised on GEF significantly outlived *ilp2-3*, *5* mutants raised on HF (Figure 3j). Thus, although GEF has effects on the expression of two insulin peptide genes, we did not detect GEF-dependent effects on insulin activity, or on the survival of insulin-deficient flies, suggesting that GEF extends life through insulin-independent means.

### Glucose-enriched food increases expression of intestine-associated cell-cell junction genes

To determine how GEF extends longevity, we used RNA sequencing (RNA-Seq) to compare transcription in whole flies raised on GEF to flies raised on HF. When we looked at differential gene expression, we found 488 upregulated genes and 555 downregulated genes in GEF-fed flies compared to HF-fed controls (Figure 4a). Gene ontology analysis of downregulated processes showed that GEF primarily leads to a decline in the expression of genes required for metabolism, and energy use (Figure 4b). In particular, we noticed significant decreases in the expression of genes involved in gluconeogenesis and lipid catabolism (Figure 4b), likely a result of the increased availability of glucose as an energy source, and consistent with our observation that flies raised on GEF have elevated triglyceride stores relative to HF-treated counterparts (Figure 1d, h).

**Figure 4.**
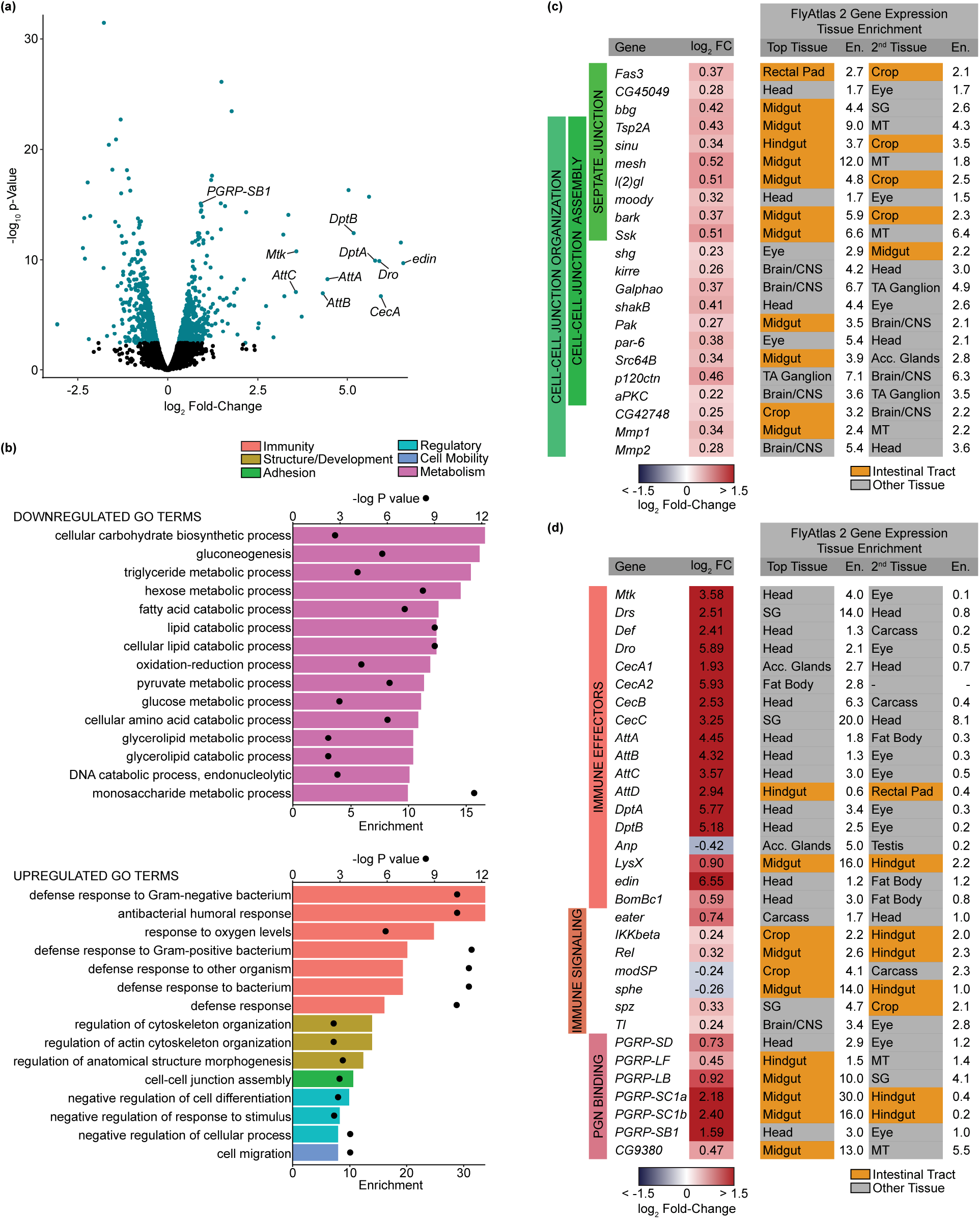
Glucose-enriched food increases expression of intestine-associated cell-cell junction genes. **(a)** Volcano plot of differentially expressed genes from comparison of flies raised on glucose-enriched food (GEF) versus unmodified holidic food (HF). Each dot represents a single gene. Teal indicates p < 0.01, FDR < 0.05. **(b)** Gene Ontology (GO) analysis from down- or up-regulated differentially expressed genes from comparison of flies raised on GEF versus HF. Bars (bottom x axis) represent enrichment scores and black circles (top x axis) represent -logP values for each enriched GO term. **(c-d)** Differentially expressed (p < 0.05) (c) cell junction genes or (d) immune-related genes from comparison of flies raised on GEF versus HF. Tissue enrichment is shown for tissues with the first and second highest enrichment scores based on FlyAtlas2 output of these genes.

In contrast to the dominance of metabolic terms among downregulated gene ontologies, we found that GEF enhanced the expression of genes involved in a number of distinct cellular processes, including immunity, cell adhesion, and cell mobility (Figure 4a, b). In fact, many of the genes with the highest GEF-dependent changes in gene expression encode antimicrobial peptides such as *attacins* and *diptericins* (Figure 4a, d). While the upregulation of immune gene expression may appear unexpected, we observed a similar phenomenon in microarray analysis comparing 10-day old female flies raised on 100 g/L glucose-enriched HF compared to females raised on unmodified HF (Figure S2). Within the list of enriched gene ontology terms for upregulated genes, we were struck by increased expression of several genes associated with cell-cell junctions (Figure 4b, c). Cell-cell junctions are critical for maintenance of epithelial structures, particularly in the intestinal tract, where barrier damage is linked to mortality (Rera et al. 2012). When we used the online resource FlyAtlas 2 to identify tissues that prominently express GEF-responsive cell-cell junction genes, we noted that a substantial number of these genes are highly expressed in the intestinal tract (Figure 4c). To confirm this, we compared transcription of representative cell-cell junction genes in whole flies, dissected heads as a control tissue, and dissected intestines. For all genes, we noted enriched expression in the intestinal tract relative to whole flies, or dissected heads (Figure S3), raising the possibility that GEF impacts organization of the gut epithelial barrier.

### Glucose-enriched food improves intestinal barrier integrity

Intestinal barrier integrity deteriorates with age and a weakened barrier is associated with reduced lifespan (Rera et al. 2012). As we observed increased expression of cell-cell junction genes in GEF-treated flies, we asked if GEF extends lifespan by improving barrier integrity.

In the fly gut, the epithelial barrier is maintained by septate junctions, which are analogous to mammalian tight junctions. Coracle (Cora), a *Drosophila* protein 4.1 homolog, is an essential component of septate junctions. As flies age, Cora and other septate junction proteins partially lose their cell membrane localization and accumulate in the cytosol, leading to breaches in the barrier, paracellular leak of lumenal material into interstitial tissue, and ultimately, death (Resnik-Docampo et al. 2017; Rera et al. 2012). To determine the effects of GEF on the intestinal barrier, we used immunofluorescence to examine the cellular distribution of Cora in the intestines of 40-day old flies raised on HF or GEF compared to 5-day old flies raised on HF. The intestines of 5-day old flies raised on HF contained orderly arrangements of large, polyploid nuclei of differentiated absorptive enterocytes, and smaller, evenly spaced nuclei of progenitor cells or secretory enteroendocrine cells (Figure 5a, Hoechst). At this young age, septate junctions are easily identified as fine margins of Cora staining at cell junctions (Figure 5a, Coracle). In 40-day old flies raised on HF, we noted classic hallmarks of age-dependent epithelial degeneration. Specifically, we detected unevenly distributed, large enterocyte nuclei, interspersed by irregular populations of smaller nuclei from progenitor/enteroendocrine cells (Figure 5a, Hoechst). In addition, we detected cytosolic accumulations of Cora (Figure 5a, asterisk), including enrichments in punctae (Figure 5a, arrowhead). In contrast, age-matched intestines of flies raised on GEF looked more similar to younger flies raised on HF, with regularly spaced nuclei (Figure 5a, Hoechst), marking cells with tight association of Cora at the plasma membrane (Figure 5a, Coracle). 3D reconstruction of 40-day old intestines highlights the difference in Cora localization between flies raised on HF or GEF (Figure 5b). In flies raised on GEF, Cora retained a reticulated pattern associated with points of cell-cell contact at septate junctions. In contrast, we detected uneven, diffuse Cora distribution in intestines of age-matched flies raised on HF.

**Figure 5.**
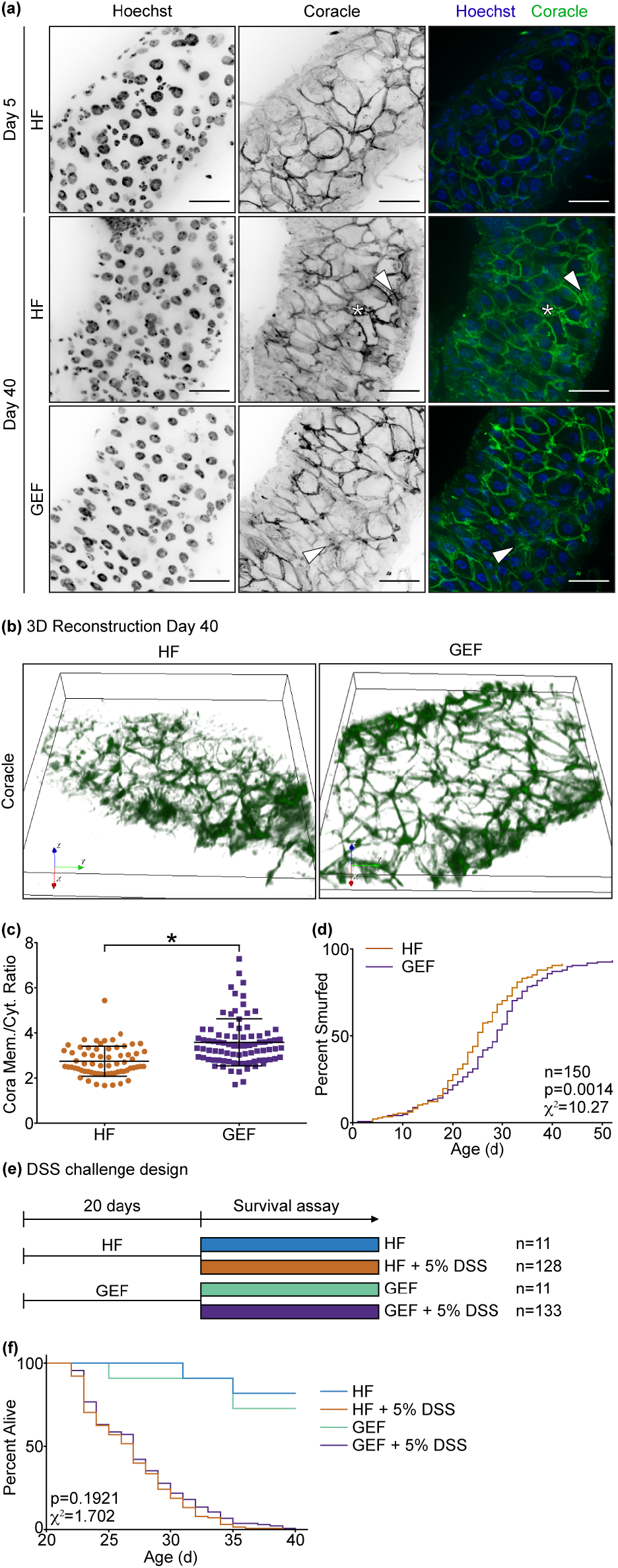
Glucose-enriched food improves intestinal barrier integrity. **(a)** Immunofluorescent images of the posterior midgut of 5- or 40-day old *w^1118^* flies raised on unmodified holidic food (HF) or glucose-enriched food (GEF) labeling DNA (Hoechst, blue) and Coracle (green). Scale bars, 25 μm. **(b)** 3D reconstruction images of Coracle in the posterior midgut of 40-day old *w^1118^* flies raised on HF or GEF. **(c)** Quantification of Coracle as a ratio of membrane to cytoplasm localization in the posterior midgut of 40-day old *w^1118^* flies raised on HF (n = 7 guts, 66 cells) or GEF (n = 8 guts, 84 cells). **(d)** Measurement of smurfs over time in *w^1118^* flies raised on HF or GEF. **(e-f)** (e) Experimental design and (f) survival curves of *w^1118^* flies raised on HF or GEF for 20 days, then transferred to food supplemented with 5% dextran sodium sulfate (DSS) or control food. Statistical significance for survival curves determined by log-rank (Mantel-Cox) test.

To quantify food-dependent impacts on the subcellular distribution of Cora, we determined the membrane to cytoplasm ratio of Cora in the midguts of flies raised on HF or GEF for 40 days. Here, we detected significantly higher membrane to cytosol ratios of Cora in 40-day old GEF-treated flies than in age-matched HF-treated flies (Figure 5c), supporting the hypothesis that GEF improves maintenance of Cora association with septate junctions as flies age.

To determine if GEF functionally improves barrier integrity in aged flies, we performed a smurf assay, in which a non-permeable dye, that only crosses the epithelium upon loss of barrier integrity, is added to the food. By counting smurfed flies over time, we found that flies raised on GEF smurfed significantly later than those on HF (Figure 5d), confirming enhanced barrier integrity in flies raised on GEF. Finally, we asked if disrupting the epithelial barrier reverts the lifespan benefits associated with GEF. For this experiment, we raised flies on GEF or HF for 20 days, at which point we transferred them to HF or GEF that we supplemented with 5% dextran sodium sulfate (DSS), a detergent that disrupts the gut barrier, for the remainder of their lives (Figure 5e). By increasing intestinal permeability with DSS, we found that flies raised on GEF completely lost their survival advantage (Figure 5f), perishing at the same time as files raised on HF. Combined, these data indicate that the lifespan extension we observe in flies raised on GEF is through a mechanism that involves maintenance of the intestinal epithelial barrier with age.

## DISCUSSION

Aging and age-related diseases pose a growing global challenge. Dietary intervention offers a promising approach to improve aging, but questions regarding optimal regimes remain. Here, we asked how glucose-enriched food (GEF) extends the lifespan of adult *Drosophila* males. We showed that GEF-dependent lifespan extension is independent of caloric restriction, or insulin activity, two established mechanisms of lifespan extension. Instead, we found that flies raised on GEF increased the expression of cell junction proteins, and had an extended period of intestinal barrier function. Furthermore, our work showed that chemical disruption of the intestinal barrier removed the lifespan extension associated with GEF-treatment. These data suggest that GEF prolongs adult viability by maintaining the integrity of the intestinal barrier for a longer period.

Maintenance of the epithelial barrier is essential for health and longevity. Occluding junctions, known as tight junctions in vertebrates or the related septate junctions of invertebrates, form this barrier, allowing for regulated movement across the epithelium (Zihni et al. 2016). Disruptions to the expression and localization of tight junction components are observed in both Crohn’s disease (Zeissig et al. 2007) and sepsis (Yoseph et al. 2016), with the upregulation of pore-forming claudin-2 and downregulation of sealing claudin-5 in both cases. Intestinal permeability also increases with age, as occluding junction proteins are downregulated (Parrish 2017). In flies, formation and maintenance of septate junction protein complexes relies on several proteins including Mesh (Izumi et al. 2012), Snakeskin (Ssk) (Yanagihashi et al. 2012), and Coracle (Lamb et al. 1998). Similar to vertebrates, disruption of septate junctions affects both health and longevity. For example, loss of Ssk affects composition of the gut bacterial community, and upregulation of Ssk extends lifespan (Salazar et al. 2018).

Effects of GEF on the intestinal barrier are consistent with previous literature that linked food intake to intestinal permeability, frequently by targeting occluding junctions (De Santis et al. 2015). For example, the amino acid glutamine has received interest for its therapeutic potential in intestinal health, as glutamine appears to directly and indirectly upregulate levels of tight junction proteins (Wang et al. 2015). Conversely, gliadin, a component of wheat, increases intestinal permeability in celiac disease through disassembly of tight junctions (Schumann et al. 2017). Gliadin binds CXCR3, inducing a MyD88-dependent release of zonulin. Loss of zonulin weakens tight junctions by altering the localization of junction proteins (Lammers et al. 2008). With their analogous role and many conserved proteins, studying the septate junctions of *Drosophila* will provide a useful *in vivo* model to explore relationships between food and the integrity of occluding junctions.

Although we did not identify the molecular mechanism by which GEF improves intestinal barrier integrity, others have explored the effect of glucose on epithelial barriers. Exposure of human retinal pigment epithelial cells to high glucose (25 mM compared to 5.5 mM) improved barrier function by increased expression of tight junction proteins (Villarroel et al. 2009). Conversely, hyperglycemia in mice, induced by streptozotocin treatment, drives intestinal barrier dysfunction by global transcriptional reprogramming of intestinal epithelial cells, specifically by downregulation of N-glycan biosynthesis genes (Thaiss et al. 2018), a critical pathway for tight junction assembly (Nita-Lazar et al. 2010). While the effect of glucose on barrier integrity is unclear, evidence suggests that glucose transporters may colocalize or interact with tight junction proteins (Rajasekaran et al. 2008). A recent study in flies suggests that dietary restriction through reduced yeast enhances barrier function via Myc activity in intestinal enterocytes (Akagi et al. 2018). Though our study was not designed to limit protein intake, our CAFE assay data indicates that flies raised on GEF received 14% of their calories from protein, whereas flies raised on HF received 38% of their calories from protein. Our RNA-seq data did not uncover differential expression of the *myc* gene. However, we cannot exclude the possibility that GEF may improve barrier integrity in a Myc-dependent manner. Future studies will be required to determine the role of Myc in GEF-dependent enhancement of the intestinal barrier.

While we focussed on cell junction genes in this report, we also observed a striking increase in expression of immune-related genes, particularly antimicrobial peptides, in GEF-treated flies. This was unexpected, as antimicrobial peptide expression increases with age (Pletcher et al. 2002) and promotes intestinal barrier dysfunction (Rera et al. 2012). Selective breeding for long-lived flies reduces age-dependent increase in antimicrobial peptide expression (Fabian et al. 2018). Furthermore, knockdown of individual antimicrobial peptides extends lifespan (Lin et al. 2018). The effect of overexpression of antimicrobial peptides on lifespan may be context-dependent as evidence suggests either detrimental (Badinloo et al. 2018) or beneficial outcomes (Loch et al. 2017). Higher baseline antimicrobial peptide expression in the long-lived GEF-treated flies suggests that the relationship between antimicrobial peptides and lifespan may be complex. As we performed RNA-sequencing on 20-day old flies, it would be of interest to measure antimicrobial peptide expression in GEF-treated flies across their lifespan to determine changes with age.

In summary, this study shows that moderate levels of glucose can extend *Drosophila* lifespan through improved intestinal barrier integrity. In humans, the intestinal barrier deteriorates with age, as well as in chronic diseases such as inflammatory bowel disease. With population aging becoming a growing global concern, further investigation of how dietary components can help maintain intestinal barrier integrity will be essential. We believe that these findings contribute to our understanding of intestinal health and may help efforts to develop preventative measures to limit the effects of aging and disease.

## EXPERIMENTAL PROCEDURES

### *Drosophila* husbandry

Virgin male *w^1118^* flies were used for all experiments unless otherwise specified. Other fly lines used were *Df(3L)Ilp2-3*,*Ilp5^3^* and *ilp2^1^ gd2HF* (Park et al. 2014). Flies were maintained at 25°C on a 12-hour light: 12-hour dark cycle and flipped to fresh food every 2-3 days. Flies in this study were allowed to develop on BDSC cornmeal food (https://bdsc.indiana.edu/information/recipes/bloomfood.html). Upon emergence, adults were transferred to their respective holidic food. The holidic food (HF) was prepared following the published protocol and recipe using the original amino acid solution (Oaa) at 100mM biologically available nitrogen (Table S1) (Piper et al. 2014). Variants to this diet included supplementation with either 50 g/L glucose (GEF), 50 g/L casein, or 22.2 g/L lard. For starvation assays, flies were maintained on 1% agar vials.

### Macronutrient assays

Samples of 5 flies were weighed and then mashed in 125 μL TE buffer (10mM Tris, 1mM EDTA, 0.1% Triton X-100, pH 7.4). Macronutrient measurements were performed in 96-well plates using commercial kits: DC Protein Assay kit (Bio-Rad, 500-0116), Triglyceride Assay kit (Sigma, TG-5-RB), and Glucose (GO) Assay kit (Sigma, GAGO20). Colorimetric readings were obtained using a microplate spectrophotometer (Molecular Devices, SpectraMax M5).

To measure circulating sugars, hemolymph was extracted from samples of 15-20 flies. Flies were carefully pierced in the thorax with a 26G needle and placed in a filter collection tube. Tubes were centrifuged at 9000g for 5 min at 4°C yielding at least 1 μL of hemolymph. 1 μL of hemolymph was diluted 1:100 in trehalase buffer (5 mM Tris pH 6.6, 137 mM NaCl, 2.7 mM KCl), and placed in a 70°C water bath for 5 min. Each sample was split into two 50 μL aliquots, one to measure glucose and one to measure trehalose. Trehalase was prepared by diluting 3 μL porcine trehalase (1 UN) in 1 mL trehalase buffer. 50 μL of this trehalase solution was added to one aliquot of each sample while 50 μL trehalase buffer was added to the other, then samples were incubated at 37°C for 24 hours. 30 μL of samples and standards were added to a 96-well plate and glucose was measured using the Glucose Oxidase (GO) Assay kits (Sigma, GAGO20). Total circulating sugars was measured from the trehalase-treated sample, free glucose was measured from the untreated sample, and trehalose was calculated as the difference between treated and untreated samples.

### Enzyme-linked immunosorbent assay (ELISA)

To measure total and circulating ILP2 levels, the *ilp2^1^ gd2HF* fly stock and protocols were provided by Dr. Seung K. Kim (Park et al. 2014). To prepare each sample, the black posterior was removed from 10 males, and the remaining bodies were transferred to 60 μL of PBS, followed by a 10 min vortex at maximum speed. Tubes were centrifuged at 1000 g for 1 min, then 50 μL of the supernatant was transferred to a PCR tube as the circulating ILP2-FH sample. To the tubes with the remaining flies, 500 μL of PBS with 1% Triton X-100 was added, homogenized with a pestle and cordless motor (VWR 47747-370), and followed by a 5 min vortex at maximum speed. These tubes were centrifuged at maximum speed for 5 min, then 50 μL of the supernatant was transferred to a PCR tube, as the total ILP2-FH sample.

For the ELISA, we used FLAG(GS)HA peptide standards (DYKDDDDKGGGGSYPYDVPDYA amide, 2412 Da: LifeTein LLC). 1 μL of the stock peptide standards (0-10 ng/ml) was added to 50 μL of PBS or PBS with 1% Triton X-100. Wells of a Nunc Maxisorp plate (Thermo Scientific 44-2404-21) were coated with 100 μL of anti-FLAG antibody diluted in 0.2M sodium carbonate/bicarbonate buffer (pH 9.4) to 2.5 μg/mL, then the plate was incubated at 4°C overnight. The plate was washed twice with PBS with 0.2% Tween 20, then blocked with 350 μL of 2% bovine serum albumin in PBS at 4°C overnight. Anti-HA-Peroxidase, High Affinity (clone 3F10) (Roche 12013819001, 25 μg/mL) was diluted in PBS with 2% Tween at a 1:500 dilution. 5 μL of the diluted anti-HA-peroxidase was added to the PCR tubes containing 50 μL of either samples or standards, vortexed, and centrifuged briefly. Following blocking, the plate was washed three times with PBS with 0.2% Tween 20. Samples and standards were transferred to wells of the plate, the plate was sealed with adhesive sealer (BIO-RAD, MSB-1001), and then placed in a humid chamber at 4°C overnight. Samples were removed with an aspirator and the plate was washed with PBS with 0.2% Tween 20 six times. 100 μL 1-Step Ultra TMB – ELISA Substrate (Thermo Scientific 34028) was added to each well and incubated at room temperature for 30 mins. The reaction was stopped by adding 100 μL 2M sulfuric acid and absorbance was measured at 450 nm on a Spectramax M5 (Molecular Devices).

### Consumption assays

For the flyPAD assay, flies were starved for 2 hours prior to the assay. HF and GEF was prepared as described, but with agarose substituted for the agar. Prepared food was melted at 95°C and then maintained at 60°C to facilitate pouring. Individual flies were placed in each flyPAD arena using a mouth aspirator at n=32 for each sample. Eating behaviour was recorded for 1 hour.

For the capillary feeding (CAFE) assay, flies were maintained in empty vials and fed liquid food through capillary tubes. To prepare liquid food for this assay, HF and GEF were prepared as described, but without the addition of agar. Flies were fed the liquid version of their respective diets for a period of 3 days. Food consumption was measured every 24 hours, and fresh food was provided each day.

### Oral glucose tolerance test (OGTT)

Flies were starved overnight for 16 hours on 1% agar, switched to vials containing 10% glucose and 1% agar for 2 hours, and then re-starved on vials of 1% agar. Samples of 5 flies were obtained after initial starvation, after 2 hours on 10% glucose, and then at both 2 hours and 4 hours following re-starvation. Samples of 5 flies were weighed and then mashed in 125 μL TE buffer (10mM Tris, 1mM EDTA, 0.1% Triton X-100, pH 7.4). Glucose was measured using the Glucose Oxidase (GO) Assay kits (Sigma, GAGO20).

### RNA isolation and RT-qPCR

To isolate RNA for both RT-qPCR and RNA-seq, samples of 5 whole flies were homogenized in 250 μL TRIzol, then incubated at room temperature for 5 min. Samples were centrifuged at 12000 g for 10 min at 4°C. Clear homogenate was transferred to a 1.5 mL Eppendorf tube, then 50 μL of chloroform was added, shaken vigorously for 15 seconds, and incubated at room temperature for 3 min. Samples were centrifuged at 12000 g for 15 min at 4°C. The upper aqueous layer was transferred to a 1.5 mL Eppendorf tube, 125 μL isopropanol was added, then left at −20°C overnight. Samples were centrifuged at 12000 g for 10 min at 4°C. The RNA pellet was washed with 500 μL 75% ethanol, centrifuged at 7500 g for 5 min at 4°C, then allowed to air dry. The RNA pellet was dissolved in RNAse free water, then incubated at 37°C for 30 min with 1 μL DNAse.

For RT-qPCR, the following primers were used in this study: *ilp2* (forward [F]: 5’-TCC ACA GTG AAG TTG GCC C-3’, reverse [R]: 5’-AGA TAA TCG CGT CGA CCA GG-3’), *ilp3* (F: 5’-AGA GAA CTT TGG ACC CCG TGA A-3’, R: 5’-TGA ACC GAA CTA TCA CTC AAC AGT CT-3’), *ilp5* (F: 5’-GAG GCA CCT TGG GCC TAT TC-3’, R: 5’-CAT GTG GTG AGA TTC GGA GCT A-3’), and *rp49* (F: 5’- AAG AAG CGC ACC AAG CAC TTC ATC-3’, R: 5’-TCT GTT GTC GAT ACC CTT GGG CTT-3’). All RT-qPCR studies were performed in triplicate, and relative expression values were calculated using delta delta Ct calculations. Expression levels were normalized to *rp49*.

### RNA-sequencing analysis

An average of 60 million reads were obtained per biological replicate. Quality check was performed with FastQC to evaluate the quality of raw, paired-end reads. Adaptors and reads of less than 36 base pairs in length were trimmed from the raw reads using Trimmomatic (version 0.36). HISAT2 ((version 2.1.0) was used to align reads to the *Drosophila* transcriptome-bdgp6, and the resulting BAM files were converted to SAM files using SAMtools (version 1.8). Converted files were counted with Rsubread (version 1.24.2) and loaded into EdgeR. In EdgeR, genes with counts less than 1 count per million were filtered and libraries normalized for size. Normalized libraries were used to identify genes that were differentially expressed between treatments. Genes with P value < 0.01 and FDR < 0.05 were defined as differentially expressed genes. Panther was used to determine Gene Ontology (GO) term enrichment of downregulated and upregulated gene sets. FlyAtlas2 was used for tissue enrichment analysis of genes of interest.

### Immunofluorescence and microscopy

Flies were briefly washed with 95% ethanol then dissected in PBS to isolate intestines. Samples were fixed for 30 min at room temperature in 4% formaldehyde. Samples were quickly washed in PBS + 0.3% Triton-X (PBT), followed by 3x 10 min washes in PBT. Samples were blocked for 1 hour in PBT + 3% bovine serum albumin (BSA) at room temperature, then incubated overnight at 4°C in PBT + 3% BSA with 1° anti-Cora 1:100 (DSHB, C615.16). Samples were washed 3x for 10 min in PBT, then incubated for 1 hour at room temperature with 2° Alexa anti-mouse 1:500. Samples were briefly washed with PBT, followed by 3x 10 min washes in PBT. Hoechst DNA stain 1:500 was added to the second 10 min wash. Samples were washed in PBS, then mounted on slides in Fluoromount (Sigma-Aldrich F4680).

Slides were visualized on a spinning disk confocal microscope (Quorum WaveFX; Quorum Technologies Inc). The R4/R5 region of the posterior midgut of each sample was located by identifying the midgut-hindgut transition and moving 1-2 frames anterior from the attachment site of the Malpighian tubules. Images were acquired using Velocity Software (Quorum Technologies). 3D reconstruction was performed with Icy.

### Quantification of Coracle

Quantification of localization of coracle in images was performed in FIJI. Three representative cells were selected per 40X image. For each cell, a transverse line was drawn across the membrane into the cell to measure coracle expression. Peak expression was recorded as the membrane value and 2.24 μm (10 px) into the cell from this peak level was recorded as the cytosol value. The membrane/cytosol ratio was calculated from these two values. This was performed in triplicate for each cell, and the average of these three measurements was recorded as the value for the cell. Sample sizes for flies raised on HF (n = 7 guts, 66 cells) and GEF (n = 8 guts, 84 cells).

### Barrier function assays

For the smurf assay, HF and GEF were prepared as described with the addition of 1% erioglaucine disodium salt (Brilliant Blue FCF). Flies were raised on their respective diets and monitored daily for extraintestinal leakage of dye or ‘smurfing’. For the dextran sulphate sodium (DSS) challenge, flies were raised on either HF or GEF for 20 days, then transferred to either HF or GEF with 5% DSS added, respectively. Deaths were recorded daily and flies were transferred to fresh food every 2-3 days.

### Statistical analysis

Statistical analysis was performed using Graphpad Prism (Version 7.0). Statistical significance was set at p < 0.05. Significance between two samples was determined by Student’s T-tests. Significance between changes over time were determined by linear regression. For lifespan and survival analysis, significance was determined using log-rank (Mantel-Cox) test. Hazard function was determined with 5-day bins.

## ACKNOWLEDGMENTS

The authors wish to thank Kin Chan at the Network Biology Collaborative Centre (nbcc.lunenfeld.ca) for the RNA-Seq service. We are grateful to Dr. Seung K. Kim for providing *ilp2^1^ gd2HF* and *Df(3L)Ilp2-3*,*Ilp5^3^* fly stocks, and sharing protocols and reagents for ILP measurement, and to Dr. Pavel M. Itskov and Dr. Carlos Ribeiro for assistance with the flyPAD. We are grateful to Dr. Andrew Simmonds for assistance in preparation of the holidic medium. Finally, we apologize to any researchers we were unable to cite due to reference limitations. This research was funded by a grant from the Canadian Institutes of Health Research to E.F. (PJT 159604) and A.G. was funded by a Natural Sciences and Engineering Research Council scholarship.

## AUTHOR CONTRIBUTIONS

The experiments were conceptualized by A.G. and E.F. Experiments were performed by A.G. Data analysis was done by A.G. and E.F. Writing of the manuscript was done by A.G. and E.F.

## DATA AVAILABILITY STATEMENT

RNA-sequencing data have been submitted to the NCBI GEO database (GSE147222).

## SUPPORTING INFORMATION LISTING

Table S1. Food composition.

Figure S1. Glucose dose longevities.

Figure S2. Microarray.

Figure S3. RT-qPCR of cell junction genes.

